# Transcription reinitiation by recycling RNA polymerase that diffuses on DNA after releasing terminated RNA

**DOI:** 10.1101/684738

**Authors:** Wooyoung Kang, Kook Sun Ha, Heesoo Uhm, Kyuhyong Park, Ja Yil Lee, Sungchul Hohng, Changwon Kang

## Abstract

Despite extensive studies on transcription mechanisms, it is unknown how termination complexes are disassembled, especially in what order the components dissociate. Our single-molecule fluorescence study unveils that RNA transcript release precedes RNA polymerase (RNAP) dissociation from DNA template in bacterial intrinsic termination of transcription much more often than concurrent dissociation. As termination is defined by release of product RNA from transcription complex, the subsequent retention of RNAP on DNA constitutes a previously unidentified stage, termed here as ‘recycling.’ During the recycling stage, RNAPs one-dimensionally diffuse on DNA in downward and upward directions, and these RNAPs can initiate transcription again at nearby promoters in case of retaining a sigma factor. The efficiency of this event, termed here as ‘reinitiation,’ increases with supplement of a sigma factor. In summary, after releasing RNA product at intrinsic termination, recycling RNAP diffuses on DNA template for reinitiation most times.

## (introduction)

Gene expression is regulated at not only initiation but also elongation and termination stages of transcription process for RNA biosynthesis, among other subsequent processes such as RNA processing and transport, and protein synthesis, processing, and targeting [1, 2]. Transcription is terminated with particular RNA sequences that are encoded by terminators in DNA template to effect particular pause of RNA polymerase (RNAP) and eventual disassembly of transcription complexes to release RNA product.

For example, in *Escherichia coli* transcription, RNA structures with a GC-rich hairpin followed by a U-rich tract cause intrinsic (factor-independent) termination, which occurs without any factors although its termination efficiency can be affected by some factors [3–6]. The U-rich tract makes elongation complex pause to allow for GC-rich RNA hairpin formation that effects conformation changes to destabilize the complex.

Although the structural changes have not been characterized yet, all termination models [7, 8] include a structural change where DNA·RNA hybrid becomes unstable, which allows RNA to get locally separated from DNA and form hairpins. Single-molecule experiments using optical tweezers have shown that termination mechanisms such as hypertranslocation and shearing mechanisms are terminator-specific [9].

However, it is still unknown yet how the complex is broken apart. Especially, which of the three components departs the transcription complex first, RNAP enzyme, RNA product, DNA template, or all together at once? To examine the temporal order of their dissociations and their post-terminational fates in *E. coli* intrinsic termination, we primarily used single-molecule fluorescence measurements.

## Results

### Construction of fluorescent transcription complex

A primary DNA template (Figure 1A) was synthesized to consist of a 50-bp upstream part including the strong promoter A1 of bacteriophage T7, a 38-bp transcription unit with the intrinsic terminator tR2 of phage λ, and a 15-bp downstream part. It was pre-labeled with Cy5 at the 5’ end of template strand (downstream end) and additionally with biotin at the 5’ end of nontemplate (coding) strand (upstream end).

**Fig. 1.**
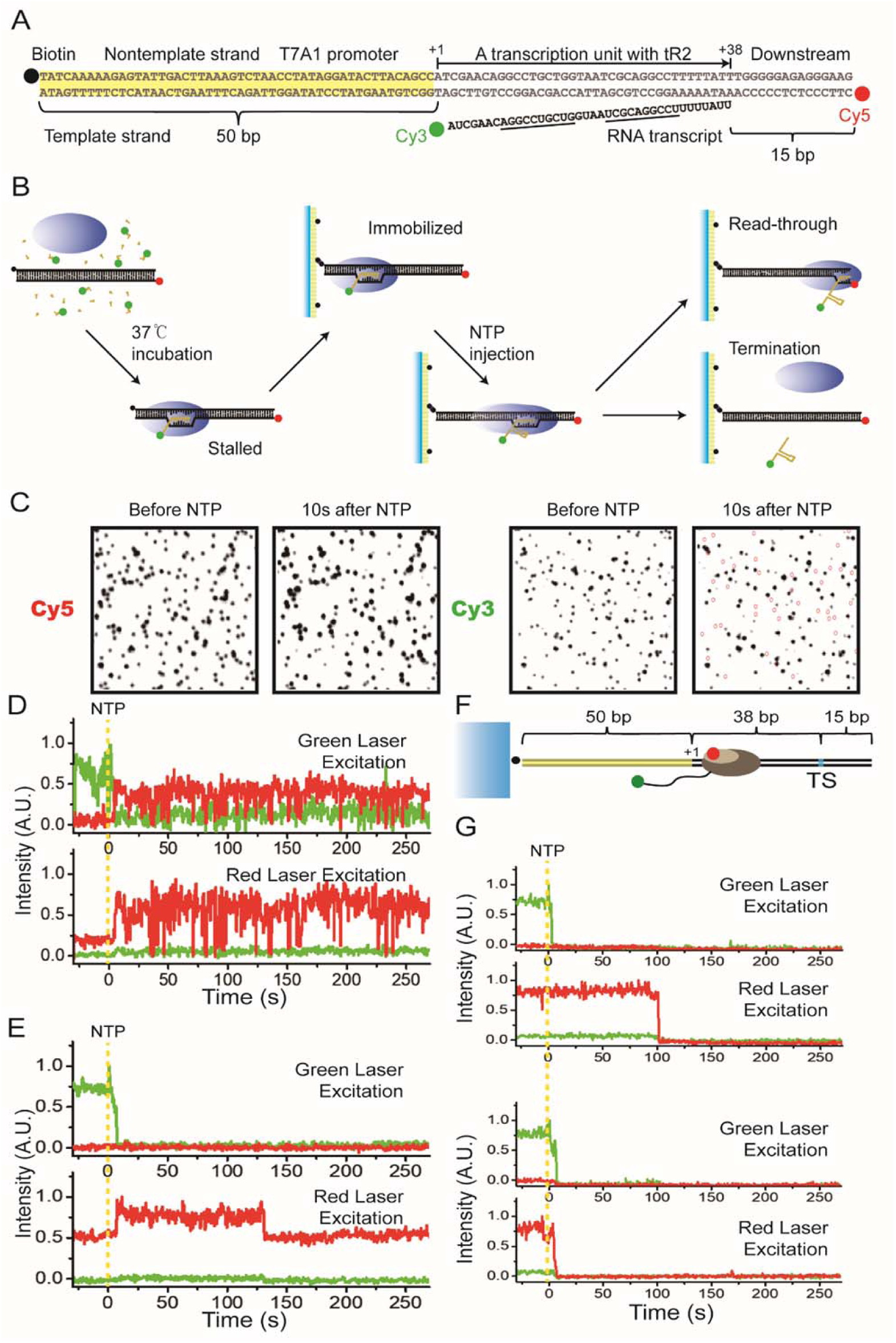
Intrinsic termination observed in single-molecule fluorescence experiments. (**A**) DNA template L+15 with T7A1 promoter and tR2 terminator was pre-labeled with biotin at the upstream end and with Cy5 at the downstream end. RNA transcript was labeled with Cy3 at the 5’-end. (**B**) Experimental scheme. Transcription complexes were stalled, immobilized and then resumed to follow readthrough or termination pathway. (**C**) Cy3-RNA and Cy5-DNA images (black) before and 10 s after NTP addition. Purple circles spot the termination complexes that have lost Cy3-RNA. (**D**-**E**) Representative fluorescence traces at green and red laser excitations from readthrough (D) and termination (E) complexes. Yellow vertical lines indicate NTP addition timing. (**F**) Construct with Cy5 placed on σ^70^ factor. (**G**) Retention of Cy5-σ on DNA after termination (top) more frequent than dissociation (bottom).

In an experiment with radioactive UTP, the major termination site (TS) was +38 position relative to the transcription start site +1, and termination efficiency was 39%. This double-labeled template with a TS-downstream length of 15 bp is denoted by L+15. Likewise, other DNA templates are denoted by L+# with TS-downstream base-pair numbers.

A fluorescent transcription complex with Cy3-labeled RNA was prepared by incubating L+15 with Cy3-labeled ApU, ATP, CTP, GTP, and *E. coli* RNAP holoenzyme with σ^70^ for 30 min in a transcription buffer (Figure 1B). Transcription is initiated preferentially with Cy3-ApU, which is incorporated into the +1 and +2 positions of RNA, but stalled at +12 waiting for the missing UTP. The stalled complexes were immobilized on polymer-coated quartz slides using biotin-streptavidin conjugation.

### Fluorescent detection of termination and readthrough

The synchronized, immobilized complexes were then subjected to resumption of elongation by adding all four ribonucleotides (NTPs), while fluorescence images of Cy3-RNA and Cy5-DNA in individual complexes were monitored using total internal reflection fluorescence microscopy. The resumed elongation is followed by either readthrough or termination at TS leading to continued or discontinued downstream transcription, respectively. In the fluorescence images (Figure 1C), vanishing of Cy3-RNA images from the Cy5-DNA spots after NTP addition indicates release of RNA from immobilized DNA, which defines termination of transcription.

In readthrough events, the terminator is ignored by RNAP that passes through TS and continues to transcribe its downstream. With readthrough complexes, from which Cy3-RNA signal is not vanished, protein-induced fluorescence enhancement (PIFE) of Cy5 occurs with red laser excitation (Figure 1D), indicating that RNAP reaches the downstream end of DNA and contacts Cy5 there [10]. Additionally, fluorescence resonance energy transfer from Cy3 to Cy5 occurs with green laser excitation, indicating that the 5’ end (Cy3) of RNA also approaches the downstream end (Cy5) of DNA presumably due to cotranscriptional folding of RNA.

In termination events, the terminator is recognized by RNAP that ends elongation at TS and does not transcribe its downstream. From termination complexes, Cy3-RNA signal vanishes 4.3±0.3 s after NTP addition (Figure 1E). The termination efficiency measured by frequency of this pathway is 34±4%, similar to the above measured using radioactive incorporation. When ITP is used instead of GTP to destabilize terminator hairpin RNA, this pathway is reduced to 10±1%. Furthermore, this pathway is not observed with a terminator-lacking template L+15M.

### RNAP remaining on DNA after releasing RNA

Surprisingly, Cy5 PIFE is observed in most (91%) of the termination complexes (Figure 1E). The PIFE starts almost concurrently with Cy3-RNA vanishing and is maintained for 536±82 s (Figure S1), while Cy5 photobleaching takes a much longer time of 2350 s (Figure S2). Accordingly, after termination, most RNAPs keep contacting DNA, as its possibility was previously speculated [11]. This was additionally observed with other intrinsic terminators (Table S3) such as *E. coli his* operon attenuator (87%) and phage ϕ82 t500 terminator (70%), suggesting that the post-terminational retention of RNAP on DNA is general.

To gain further support, we repeated the experiments with Cy5 placed on σ factor rather than DNA (Figure 1F), because the initiation factor has been recently reported to remain in some elongation complexes [12, 13]. *E. coli* σ^70^ was labeled with Cy5 at Cys-336 as previously described [14]. Cy5-σ is retained in 75% of the initially-stalled complexes (Figure S3), which falls within a previously reported 70-90% range of σ retention [13].

This Cy5-σ of holoenzymes mostly (84%) remain on immobilized DNA even after Cy3-RNA vanishing of termination (Figure 1G). In no cases, Cy5-σ vanishes before Cy3-RNA. The dissociation time of σ^70^ (588±85 s, Figure S4) is similar to that of RNAP (536±82 s, Figure S1), suggesting that the holoenzyme maintains a stable complex throughout its retention on DNA.

### One-dimensional diffusion of RNA-free RNAP on DNA

After releasing RNA, RNAP can reach the Cy5-end of DNA through three-dimensional (3D) diffusion after dissociation from DNA or through one-dimensional (1D) diffusion (sliding or hopping) without dissociation from DNA. To examine how RNA-free RNAP reaches DNA ends, a recognition sequence of EcoRI restriction enzyme was inserted into DNA at downstream of TS (Figure 2A). DNA-bound EcoRI blocks procession of *E. coli* RNAP [15], so this roadblock would obstruct 1D diffusion downward from TS, but not TS-upward 1D diffusion or 3D diffusion.

**Fig. 2.**
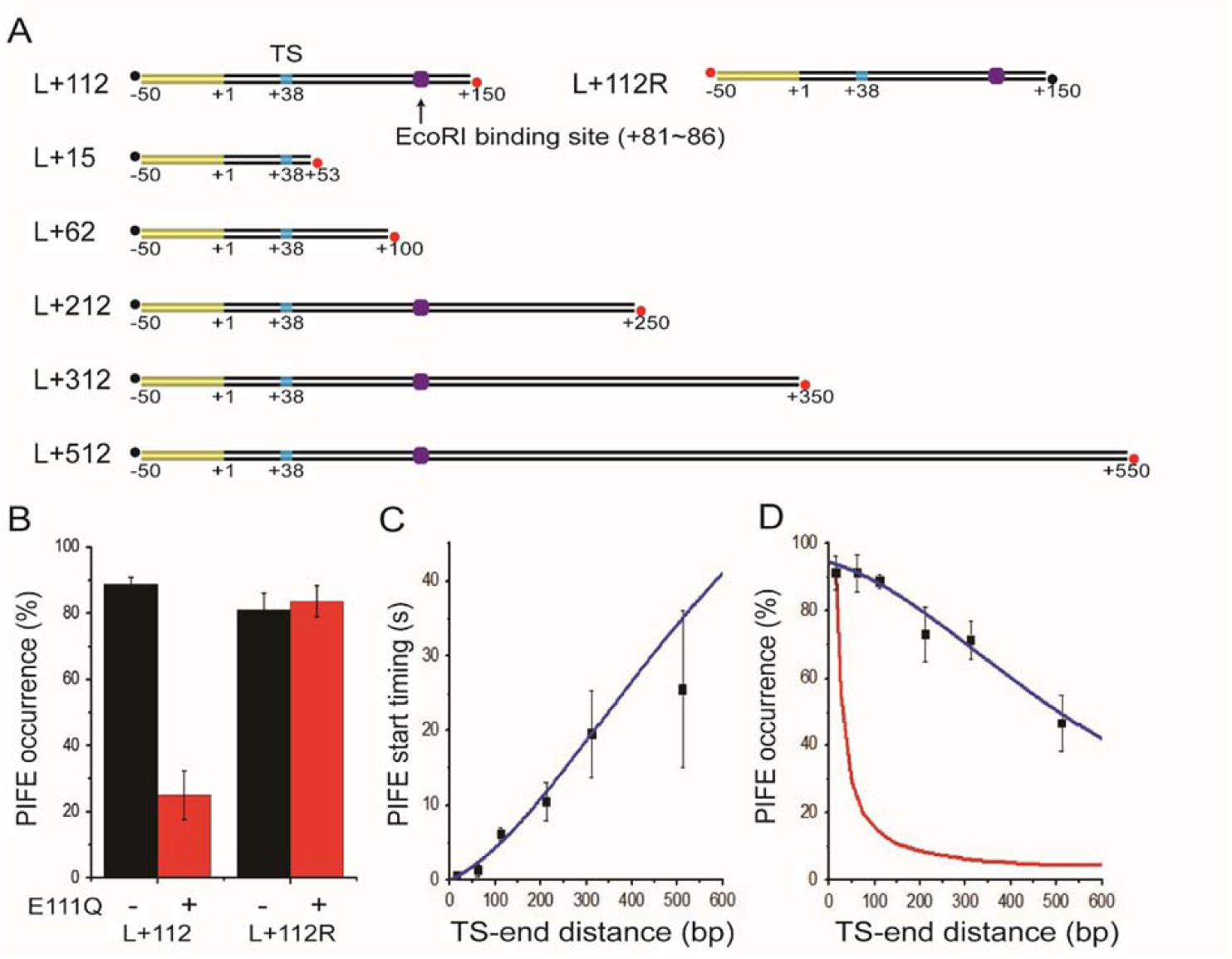
Diffusion of RNAP on DNA after termination. (**A**) DNA templates with varying distance between TS (cyan) and Cy5-labeled end (red) with or without an EcoRI binding site (purple). (**B**) Reduction of Cy5 PIFE occurrence by a roadblock on L+112 but not L+112R. (**C**) PIFE start timings plotted against TS-end distances. Cy5 PIFE start was timed since Cy3 vanishing from six templates, L+15 through L+512 (black). The timing data fit to a 1D diffusion model (blue). (**D**) PIFE occurrences plotted against TS-end distances. Cy5 PIFE was counted with the six templates (black). The occurrence data fit to the same 1D diffusion model (blue) but not a 3D diffusion model (red).

In experiments using a cleavage-lacking mutant E111Q of EcoRI [16], this roadblock reduces PIFE occurrence among the termination complexes when Cy5 is placed at the downstream end of L+112 (from 89±2% to 25±7%), but not with Cy5 placed at the upstream end of L+112R (Figures 2A and 2B). As the PIFE occurrence reduction is as much as DNA cleavage inhibition by E111Q (71±5%; Figure S5), the Cy5-ends are reached by RNAP mostly through 1D diffusion, much more often than 3D diffusion.

### Post-terminational diffusion lifetime of RNAPs

It is known that RNAP preferentially binds blunt ends of DNA, so the post-terminational retention time of RNAP or σ on short DNA molecules could be longer than the 1D diffusion lifetime, which can be measured on long DNA molecules where diffusing RNAP takes too long to reach an end. The σ-retention time (Cy5 signal duration) after termination was measured on a long DNA of 1560 bp with 6.4% of PIFE occurrence expectation.

The post-terminational diffusion of RNAP lasts for 75±5 s on this long DNA (Figure S6). This lifetime is clearly shorter than RNAP’s association with a short DNA of 103 bp at Cy5- end (Cy5-PIFE duration of 536±82 s; Figure S1) or similar association of Cy5-σ holoenzyme with the short DNA (Cy5 signal duration of 588±85 s; Figure S4). RNAP is thus apparently trapped for an extended period once it reaches the ends of DNA.

### Post-terminational diffusion coefficient of RNAPs

The 1D diffusion coefficient of post-terminational RNAPs can be estimated by the correlations that Cy5 PIFE starts later and occurs less frequently as the Cy5-end is placed further away from TS (Figures 2C and 2D). Cy5 PIFE was measured with six templates, L+15, L+62, L+112, L+212, L+312, and L+512, where the distance between TS and Cy5-end of DNA (TS-downstream length) varies from 15 to 512 bp (Figure 2A).

Termination efficiency (ranging from 31±3% to 36±16%) or timing (ranging from 4.0±0.8 to 7.1±1.3 s) is little affected by the TS-downstream length (Table S1; Figure S7). On the other hand, with increasing TS-end distance, the PIFE start delay increases from 1.8±0.5 to 30±19 s since RNA release (Figure 2C), and the PIFE occurrence decreases from 91% to 47% (Figure 2D), both supporting for 1D diffusion of RNAP on DNA after termination.

RNAP diffusion on DNA is modeled to occur with one reflecting end on the surface side and the other absorbing end on the buffer side, which is reasonable because PIFE occurrence at free ends is highly probable with short DNA molecules. This 1D diffusion model adopting the above-determined post-terminational diffusion lifetime of RNAPs fits well to the correlation of the PIFE start timing (*R*^2^ = 0.94) or occurrence (*R*^2^ = 0.96) with the TS-end distance (Figure 2C and 2D), from which the diffusion coefficient is estimated as 3.5×10^−4^ or 3.4×10^−4^ μm^2^/s, respectively (see the STAR Methods).

On the other hand, the PIFE occurrence data do not fit to a previously described 3D diffusion model [17] (Figure 2D). The PIFE occurrence at zero distance can be additionally estimated by the above 1D diffusion model, and 94% of complexes are estimated to retain RNAP at the time of RNA release at TS, and the remaining 6% release RNAP together with RNA.

### Transcription reinitiation by post-terminational RNAPs

During post-terminational diffusion on DNA, RNA-free holoenzymes of RNAP could initiate transcription again when they encounter a promoter. Even core enzymes of RNAP could do so if they become adequately complemented with σ. Here ‘reinitiation’ refers to another round of promoter binding and initiation by the RNAP that has released RNA but not fallen off DNA and diffuses on DNA rather than by 3D association of RNAP.

Reinitiation was examined using a template with two transcription units (Figure 3A). L+15 containing a short transcription unit was extended downstream to contain a roadblock site and another long transcription unit, among other extensions. As explained above, short transcripts from the upstream unit can be labeled with Cy3-ApU, which reports whether termination or readthrough occurs at TS. Long transcripts from the downstream unit contain five repeating 21-bp sequences, and can be probed by a Cy5-labeled DNA oligomer complementary to the repeat sequence.

**Fig. 3.**
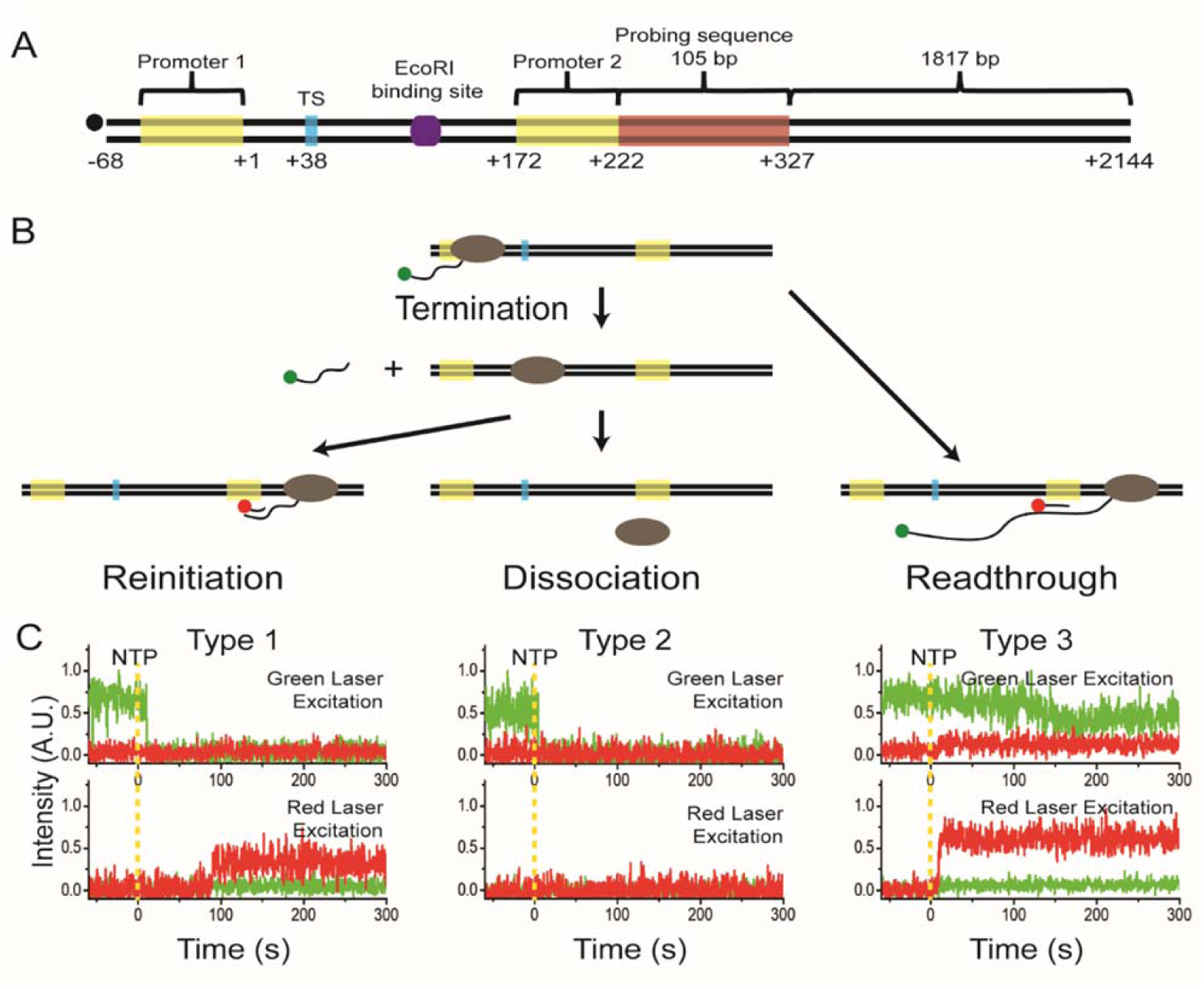
Detection of transcriptional reinitiation. (**A**) A DNA template contains two transcription units with a roadblock (purple) between them. Transcripts from the downstream unit have a probing target (brown). (**B**) Three possible courses are reinitiation on the downstream unit after TS termination, dissociation of RNAP after TS termination, and readthrough at TS with continued downstream transcription. (**C**) Three types of fluorescence time-traces were observed; Cy3 vanishing with Cy5 probing (type 1), Cy3 vanishing without Cy5 probing (type 2), and Cy3 nonvanishing with Cy5 probing (type 3).

Three courses can proceed (Figure 3B) as they were observed in different types of fluorescence time-traces (Figure 3C). First, reinitiation on the downstream unit occurs after termination from the upstream unit, as Cy5-oligomer probing signal follows Cy3-RNA signal vanishing (type 1). Second, dissociation of RNAP occurs after TS termination, as no Cy5 probing follows Cy3 vanishing (type 2). Third, readthrough occurs for continued downstream transcription, as Cy5 probing follows Cy3 nonvanishing (type 3). The relative frequencies of types 1, 2, and 3 were 0.10±0.05, 0.41±0.10, and 0.49±0.12, respectively.

Because transcript probing was not complete, not all readthrough events were counted, and some reinitiation events were observed as type 2 instead of type 1. However, using the measured termination efficiency (34%), the probing efficiency was estimated as 49%, allowing the reinitiation (0.21) and dissociation (0.30) events to be re-sorted as described in the Methods. Thus, the reinitiation portion among the termination events was 41% with a mixture of holoenzymes and core enzymes. The reinitiation efficiency decreases to 19% with the above-mentioned roadblock, but increases to 59% with addition of 3 μM σ^70^.

## Discussion

This study discovered that RNA transcript is released first from transcription complex, and RNAP dissociates from DNA template much later most times in *E. coli* intrinsic termination, based on our single-molecule monitoring of fluorescent RNA transcript, DNA template or σ factor. This sequential release (94%) is much more frequent than their concurrent dissociation (6%). Considering that only 75% of elongation complexes retained σ factor in our case, the post-terminational retention of RNAP on DNA seems to occur similarly with both core and holoenzymes. Additionally, we showed that most holoenzymes retain the σ factor after termination.

Because termination should refer to RNA release only, the subsequent retention of RNAP on DNA constitutes a previously unidentified stage. This fourth stage of transcription after the initiation, elongation, and termination stages is here termed as ‘recycling’ (Figure 4). During the recycling stage, post-terminational RNAPs apparently diffuse on DNA one-dimensionally in downward and upward directions.

**Fig. 4.**
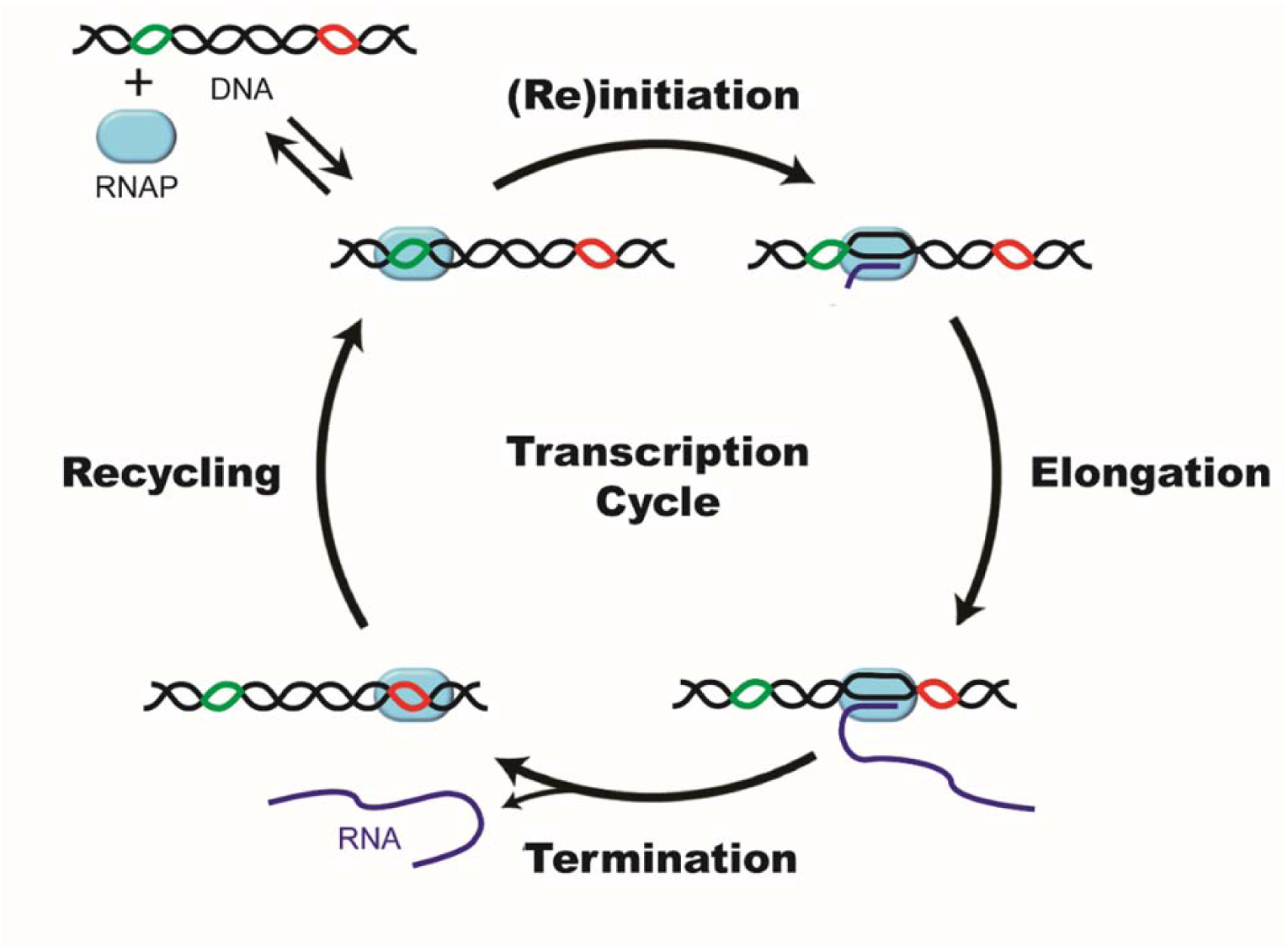
Four stages of transcription cycle. (1) Initiation: After binding DNA (black) at a promoter (green), RNAP (cyan) incorporates several NTPs into RNA (blue), and advances downward clearing the promoter. (2) Elongation: RNAP further advances processively downward, resuming from occasional pause or backtrack. (3) Termination: RNAP pauses at a terminator (red) to undergo conformational changes, and releases RNA product. (4) Recycling: RNAP diffuses on DNA downward and upward until it reinitiates transcription at the same or another promoter unless RNAP dissociates off DNA.

One-dimensional diffusion of RNAP has been suggested to accelerate promoter search [18-20], but few experiments have directly proved it. Furthermore, RNAP diffusion on DNA was recently measured to be too short (<30 ms) to contribute to initiation kinetics under physiological conditions [21]. This lifetime of pre-initiational RNAP on DNA is much shorter than that of post-terminational RNAP (>1 min) observed in this study, suggesting that their conformations are distinct from each other. Our single-molecule detection of such long and frequent (95%) retention of post-terminational RNAP on DNA provides the first direct evidence that 1D diffusion of RNAP can dramatically accelerate the promoter search process at least *in vitro*.

RNAP’s post-terminational retention of σ and diffusion on DNA allow for transcription reinitiation, which is defined to occur by promoter binding through 1D rather than 3D diffusion of RNAP. Reinitiation is demonstrated here to occur on a downstream promoter oriented in the same direction with the upstream promoter. Although it is not known yet whether 1D-diffusing RNAP can flip to reinitiate at an oppositely oriented promoter, RNAP can diffuse backward or upward towards the upstream end of DNA. Furthermore, σ supplement is shown here to increase the reinitiation efficiency, suggesting that even core enzyme can become reinitiation-competent during the recycling stage, which ends with reinitiation or dissociation of RNAP.

Post-terminational retention and diffusion of RNAP on DNA and transcription reinitiation competence of such RNAP during the recycling stage have several functional implications, although they have not been observed to occur *in vivo* yet. First, they would facilitate transcriptional burst, which has been observed for both bacterial and eukaryotic transcriptions [22-24]. Post-terminational retention of RNAP on DNA during the recycling stage may increase its local density and render it readily convertible into reinitiation-competent state, contributing to the transcriptional burst.

Second, simultaneous transcriptional regulation of multiple operons can be facilitated. In bacterial genomes, functionally related genes and operons are often clustered and regulated together as a regulon [25-28]. One can hypothesize that a single RNAP molecule continually transcribes clustered operons while diffusing on inter-operon regions of DNA chromosome.

Third, transcriptional memory can be facilitated by recycling RNAPs. A loop formed between promoter and terminator is observed in some eukaryotic genes, although not yet in bacterial operons. Such loops can increase initiation efficiency and provide transcriptional memory [29-31], where facilitated diffusion of RNAPs on the loop could speed up reactivation.

In this work, we established a single-molecule fluorescence assay to study bacterial transcription termination dynamics. Our construction of fluorescent transcription complexes can permit single-molecule monitoring of their components in studies on transcriptional elongation, termination, recycling, and reinitiation.

## Methods

### Preparation of transcription templates

All transcription templates were prepared by polymerase chain reactions using AccuPower ProFi Taq PCR premix from Bioneer, Korea with the amplification templates and primers purchased from Integrated DNA Technologies, USA. Their sequences are listed in table S3. All amplification products were purified using the Cleanup kit Expin PCR SV mini from GeneAll, Korea.

DNA template L+15 was prepared using UP8_template, 5’ biotin-labeled forward_primer_ biotin, and 5’ Cy5-labeled reverse_primer_L+15; L+15M using UP8M_template, 5’ biotin-labeled forward_primer_biotin, and 5’ Cy5-labeled reverse_primer_L+15M; L+62 using UP8_template, 5’ biotin-labeled forward_primer_biotin, and 5’ Cy5-labeled reverse_primer_L+62; L+112 using UP8_template_2, 5’ biotin-labeled forward_primer_biotin, and 5’ Cy5-labeled reverse_primer_ L+112; and L+112R using forward_primer_Cy5’ and reverse_primer_biotin instead. Template for phage ϕ82 t500 terminator was prepared using t500_template, 5’ biotin-labeled forward_ primer_biotin, and 5’ Cy5-labeled reverse_primer_L+62; and template for *E. coli his* operon attenuator using his_template, 5’ biotin-labeled forward_primer_biotin, and 5’ Cy5-labeled reverse_primer_L+62.

For L+212 and L+312 constructions, L+112 and additional_part_1 were annealed with DNA_splint_1 by cooling from 90°C to 30°C for 120 min in the annealing buffer (10 mM Tris-HCl, pH 8.0, with 50 mM NaCl), ligated using T4 DNA ligase 2 purchased from New England Biolabs (NEB), USA, and used for amplification reactions. L+212 was prepared using forward_primer_ biotin and reverse_ primer_L+212; L+312 using forward_primer_biotin and reverse_primer_ L+312. For L+512 construction, L+312 and additional_ part_2 were annealed with DNA_splint_2 and the same ligation reaction was repeated. L+512 was prepared using forward_primer_biotin and reverse_primer_L+512.

For templates used for reinitiation detection and σ retention time estimation, long_tail template with HindIII recognition sequence was prepared using lambda DNA (NEB), lambda_forward_primer, and lambda_reverse_primer. To construct the upstream part of reinitiation template, L+112_α was prepared with UP8 template_2, forward_primer_extension, and 5’ reverse_primer_L+112. L+112_α and reinitiation_part_1B were annealed with DNA_splint_reinitiation by cooling from 90 to 30 °C for 120 min in the annealing buffer, and the same ligation reaction was repeated. Long_tail DNA, the upstream part of reinitiation template, and L+512 were each digested with HindIII (NEB) for one h at 37 °C in the CutSmart™ buffer (NEB). After that, HindIII was deactivated for 20 min at 80 °C.

Long_tail DNA were annealed with cleaved L+512 or the upstream part of reinitiation template by the same protocol as above and used for amplification reaction. L+lambda was prepared using forward_primer_biotin and lambda_forward_primer. DNA template for reinitiation detection was prepared using forward_primer_biotin_α and lambda_reverse_ primer.

### Preparation of Cy5-labeled σ factor

An N-terminal His_6_-tagged *E. coli* σ^70^ was produced in BL21(DE3) cells at 25 °C. Soluble fractions of cell lysates were purified by a 5-ml HisTrap column (GE Healthcare) followed by a 5-ml HiTrap heparin column (GE Healthcare). Purified σ^70^ was incubated with Cy5 maleimide mono-reactive dye (1 mM, GE Healthcare) in the storage buffer (50 mM Tris-HCl, pH 8.0, 100 mM NaCl, and 0.1 mM EDTA) overnight at room temperature. Unreacted Cy5-maleimide molecules were removed using an Amicon ultra centrifugal filter (Merck).

### Single-molecule transcription termination experiments

To construct a transcription complex, we incubated DNA template (50 nM) with *E. coli* RNAP holoenzyme (20 nM, NEB), ATP, GTP, CTP (20 μM each, GE Healthcare), and Cy3-labeled ApU (250 μM, TriLink) for 30 min at 37 °C. For the experiments with Cy5-labeled σ, we used *E. coli* RNAP core enzyme (20 nM, NEB) plus 1 μM Cy5-σ instead of RNAP holoenzyme. Quartz slides were cleaned using piranha solution to remove organic residues, incubated with (3-aminopropyl)trimethoxysilane (United Chemical Technologies), and coated with polymers by incubating the slides in a 1:40 mixture of biotin-PEG-5000 and m-TEG-5000 (Laysan Bio). The slides were treated with streptavidin (0.2 mg/ml, Invitrogen) for 5 min, and transcription complexes were immobilized on the surface of quartz slides via biotin-streptavidin conjugation.

### Single-molecule fluorescence imaging

A home-made total-internal-reflection fluorescence microscope is equipped with a 532-nm green laser (EXLSR-532-50-CDRH, Spectra-physic) for Cy3 excitation and with a 640-nm red laser (EXLSR-635C-60, Spectra-physic) for Cy5 excitation. An electron multiplying charge-coupled device camera (Ixon DV897, Andor Technology) was used as an imaging device and controlled by using a customized C# program. All experiments were performed at 37 °C with 0.05 s exposure time. The time resolution of the experiments was 0.1 s because we used the alternating laser excitation (ALEX) mode.

All experiments were performed with an oxygen scavenging buffer (10 mM Tris-HCl, pH 8.0, 20 mM NaCl, 20 mM MgCl_2_, 1 mM dithiothreitol, 5 mM 3,4-protocatechuic acid, and 100 nM protocatechuate-3,4-dioxygenase) and 200 μM NTP each. In the roadblock experiments, the immobilized transcription complexes were incubated with E111Q (1 nM) for 10 min, and DNA templates without E111Q were digested using wild-type EcoRI (10 units/μl, Beams Biotechnology). Then, we performed the experiments in the oxygen scavenging buffer containing 1 nM E111Q.

### Single-molecule transcription reinitiation experiments

The imaging buffers before and after NTP injection contained the same concentration of Cy5-labeled ‘imager DNA’ (50 nM, Cy5-TGTGT GTGGT CTGTG GTGTC T) to maintain the same background level. From our analysis were excluded the traces that showed Cy5 signal before NTP addition (nonspecific probing). Because tR2 termination occurs at 4.3 s after NTP addition on average, only Cy3 vanishing within 15 s was regarded as termination. Among these traces, we classified those with Cy5 probing within 240 s as type 1 and the others as type 2. On the other hand, those with Cy3 surviving longer than 180 s and Cy5 probing within 240 s were classified as type 3. Other traces were excluded from data analysis. The exclusion due to Cy3 survival within the range of 15 to 180 s was minor (11%). The exposure time of 0.2 s was used to reduce photobleaching.

### One-dimensional diffusion model

Diffusion of RNAP on DNA can be modeled as 1D diffusion of a point particle with one reflecting end and one absorbing end. RNAP diffusion coefficient and RNAP retention on DNA such as PIFE occurrence and start timing can be estimated using this model

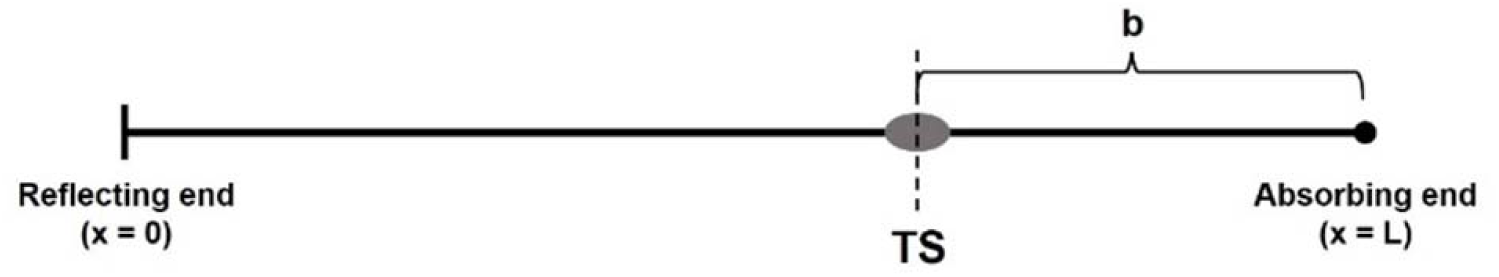

The mean time to reach the absorbing end, *W*(*x*), is defined as a function of the distance between the reflecting end and termination site (TS), *x*. Then, *W*(*x*) satisfies the equation (1).

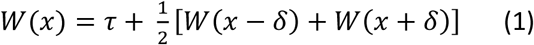

where δ is a step size of the random walk of the particle moving rightward or leftward every τ s. Because RNAP does not remain on DNA permanently, an RNAP dissociation term is added to the equation (1).

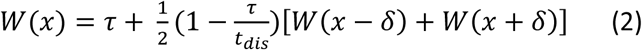

where *t*_*dis*_ is a dissociation time of RNAP after termination. In the limit δ → 0, we can get the differential equation (3) for *W*(*x*).

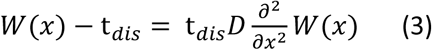

where D is a diffusion coefficient, defined as δ^2^/2τ.

When TS is positioned at the reflecting end (*x* = 0), the mean capture time does not change with *x*. When TS is positioned at the absorbing end (*x* = L), the particle is captured right after the particle release, generating the boundary conditions (4) and (5).

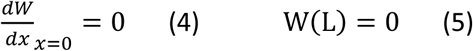

With these boundary conditions, the equation (3) is solved as a function of *x* and the total length *L* as shown in equation (6).

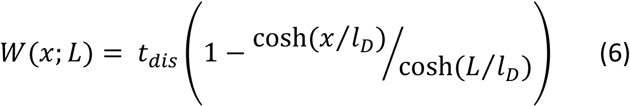

where *l*_*D*_^2^ is equal to *t*_*dis*_D. Since *x* is fixed as 88 bp in our experiments, we can rewrite the equation (6) as a function of the distance between TS and the absorbing end, denoted by b.

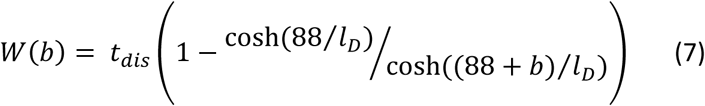

Diffusion coefficient of RNAP can be estimated by fitting PIFE timing data to the equation (7).

Similarly, the probability of capture on the absorbing end, *P*(*x*), is defined as a function of x. Then, *P*(*x*) also satisfies the equation (8).

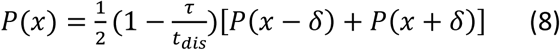

In the limit δ → 0, we can get the differential equation (9) for *P*(*x*).

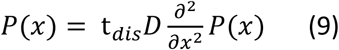

When TS is positioned at the reflecting end, the probability of capture also does not change with x. Otherwise, the probability of capture is equal to the RNAP retention probability during termination at TS when TS is positioned at the absorbing end. These two conditions generate two boundary conditions (10) and (11).

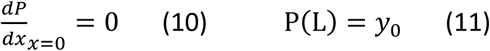

where y_0_ is the RNAP retention probability during termination at TS. With these boundary conditions, we can solve the equation (9) to get the equation (12).

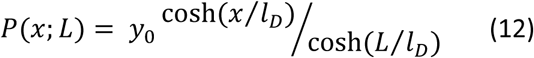

We can rewrite the *P*(*x*) as a function of b.

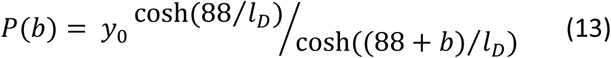

RNAP retention probability during termination at TS can be estimated by fitting PIFE occurrence to the equation (13).

### Calculation of reinitiation efficiencies

Among the three types of fluorescence traces of Fig. 3C, type 3 uniquely represents TS-readthrough events (Fig. 3B), but does not cover all of them because transcript probing is not complete. The unprobed readthrough events are not distinguishable from non-activated events and could not be directly counted.

On the other hand, types 1 (0.101) and 2 (0.406) together represent total TS-termination events (Fig. 3B), but their sum (0.507) is an overestimation because of incomplete probing. However, using the measured termination efficiency of 33.5%, the unprobed readthrough events (type 4) can be estimated to be 0.513, i.e. (0.507−0.335)/0.335. Types 3 (0.493) and 4 (0.513) together represent total readthrough events (1.006), with probing efficiency of 49.0%. Then, total readthrough and termination events become 1.513.

Due to incomplete probing, some reinitiation events must have been shown as type 2 instead of type 1. Assuming that probing efficiency was the same in readthrough and termination events, total probed and unprobed reinitiation events can be estimated to be 0.206 (= 0.101/0.490), and the remaining dissociation events being 0.301 (= 0.507 − 0.206). Thus, the reinitiation portion among the total termination events is 40.6% (= 0.206/0.507) with a mixture of holoenzymes and core enzymes.

## Acknowledgments

We thank Prof. Robert Landick for plasmid pRPODS366C encoding a single-cysteine derivative of σ^70^. This work was supported by grants from the National Research Foundation of Korea (2019R1A2C2005209 to SH and 2017R1A2B4002213 to JYL), the High Risk High Return Project of KAIST (N10110078 to CK), and the Research Fund of UNIST (1.160106.01 to JYL).

## Author contributions

C.K. and S.H. conceived the study. K.S.H. constructed transcription complexes and performed molecular biology experiments. W.K., H.U., K.P., and J.Y.L. measured the single-molecule fluorescence images and performed biophysics experiments. C.K. supervised the molecular biology part. S.H. supervised the biophysics part. All analyzed the data and contributed to writing of the paper.

