## Supplemental Information for "Transcription reinitiation by recycling RNA polymerase that diffuses on DNA after releasing terminated RNA"

[1]Department of Physics and Astronomy, and Institute of Applied Physics, Seoul National University, Seoul 08826, Republic of Korea. [2]Department of Biological Sciences, Korea Advanced Institute of Science and Technology, Daejeon 34141, Republic of Korea. [3]School of Life Sciences, Ulsan National Institute of Science and Technology, Ulsan 44919, Republic of Korea. [4]Present address: Department of Life Science, University of Suwon, Gyeonggi-do 18323, Republic of Korea. [5]Present address: Department of Physics, University of Oxford, Oxford OX1 3PU, United Kingdom. [6]These authors contributed equally: Wooyoung Kang, Kook Sun Ha. [7]These authors jointly supervised this work: Sungchul Hohng, Changwon Kang. *Correspondence and requests for materials should be addressed to C.K. or S.H.

### Supplementary Tables

**Table S1. Termination efficiency and timing of various templates**

| Sample | Termination efficiency (%) | Termination timing (s) |
|---|---|---|
| L+15 | 34 ± 4 | 4.3 ± 0.3 |
| L+62 | 31 ± 3 | 4.2 ± 0.5 |
| L+112 | 35 ± 4 | 5.5 ± 0.7 (without EcoRI E111Q), 4.9 ± 0.7 (with EcoRI E111Q) |
| L+212 | 34 ± 1 | 4.0 ± 0.8 |
| L+312 | 36 ± 13 | 5.2 ± 1.3 |
| L+512 | 36 ± 16 | 5.9 ± 1.4 |
| L+112R | | 7.1 ± 1.3 |

**Table S2. Termination timing under various conditions in reinitiation experiments**

| Condition | Termination timing (s) |
|---|---|
| Without sigma | 5.9 ± 1.3 |
| With sigma | 6.2 ± 1.3 |
| EcoRI E111Q binding | 6.0 ± 2.3 |

### Table S3. Template DNA sequence

| Oligo name | Sequence (5′ → 3′) |
|---|---|
| UP8_template | TATCA AAAAG AGTAT TGACT TAAAG TCTAA CCTAT AGGAT ACTTA CAGCC ATCGA ACAGG CCTGC TGGTA ATCGC AGGCC TTTTT ATTTG GGGGA GAGGG AAGTC ATGAA AAAAC TAACC TTTGA AATTC GATCT CCAGG ATCCA CCACC |
| UP8M_template | TATCA AAAAG AGTAT TGACT TAAAG TCTAA CCTAT AGGAT ACTTA CAGCC ATGGC CTGCT GGTGA CTGAC TGACT GACTG AC |
| t500_template | TATCA AAAAG AGTAT TGACT TAAAG TCTAA CCTAT AGGAT ACTTA CAGCC ATCCC AAAGC CCGCC GAAAG GCGGG CTTTT CTGTT TCTGG GCGGT GAAGT CATGA AAAAA CTAAC CTTTG AAATT CGATC TCCAG GATCC ACCAC C |
| his_template | TATCA AAAAG AGTAT TGACT TAAAG TCTAA CCTAT AGGAT ACTTA CAGCC ATCCG AAAGC CCCCG GAAGA UGCAU CUUCC GGGGG CUUUU UUUUU TGGGC GGTGA AGTCA TGAAA AAACT AACCT TTGAA ATTCG ATCTC CAGGA TCCAC CACC |
| UP8_template_2 | GCGAG ATTAC CATTA AGTGA ATTCG AAAAA AGCAC GCTAC CGCCC CAGGC GGTGG TGGAT CCTGG AGATC GAATT TCAAA GGTTA GTTTT TTCAT GACTT CCCTC TCCCC CAAAT AAAAA GGCCT GCGAT TACCA GCAGG CCTGT TCGAT GGCTG TAAGT ATCCT ATAGG TTAGA CTTTA AGTCA ATACT CTTTT TGATA |
| additional_part | pAATTC TTACA ATTTA GACCC TAATA TCACA TCAGA CACTA ATTGC CTCTG CCAAA ATTCT GTCCA CAAGC GTTTT AGTTC GCCCC AGTAA AGTTG TCAAT AACGA CCACC AAATC CGCAT GTTAC GGGAC TTCTT ATTAA TTCTT TTTTC GTGGG GAGCA GCGGA TCTTA ATGGA TGGCG CCAGG TGGTA TGGAA GC |
| additional_part_2 | pGGGCT GAAAG TAGCG CCGGG TAAGG TACGC GCCTG GTATG GCAGG ACTAT GAAGC CAATA CAAAG GCTAC ATCCT CACTC GGGTG GACGG AAACG CAGAA TTATG GTTAC TTTTT GGATA CGTGA AACAT GTCCC ATGGT AGCCC AAAGA CTTGG GAGTC TATCA CCCCT AGGAC ACACA AGACA CCACA AGCTT AGACC |
| DNA_splint | TGTGA TATTA GGGTC TAAAT TGTAA GAATT GCGAG ATTAC CATTA AGTGA ATTCG AAAAA |
| DNA_splint_2 | GCGTA CCTTA CCCGG CGCTA CTTTC AGCCC GCTTC CATAC CACCT GGCGC CATCC ATTAA |
| reinitiation_part_1B | pTAATA TCACA TCATT AGACA CTTAT CAAAA AGAGT ATTGA CTTAA AGTCT AACCT ATAGG ATACT TACAG CCTGC AGACA CCACA GACCA CACAC AAGAC ACCAC AGACC ACACA CAAGA CACCA CAGAC CACAC ACAAG ACACC ACAGA CCACA CACAA GACAC CACAG ACCAC ACACA AGACA CCACA AGCTT AGACC |
| DNA_splint_reinitiation | AGTGT CTAAT GATGT GATAT TAGCG AGATT ACCAT TAAGT GAATT CGAAA AA |

### Table S4. Primer sequence

| Oligo name | Sequence (5′ → 3′) |
|---|---|
| forward_primer_biotin | Biotin-TATCA AAAAG AGTAT TGACT TAAAG TC |
| reverse_primer_biotin | Biotin-GCGAG ATTAC CATTA AGTGA A |
| forward_primer_Cy5 | Cy5-TATCA AAAAG AGTAT TGACT TAAAG TC |
| reverse_primer_L+15 | Cy5-CTTCC CTCTC CCCCA AATAA AAAG |
| reverse_primer_L+15M | Cy5-GTCAG TCAGT CAGTC AGTCA CCAGC AG |
| reverse_primer_L+62 | Cy5-GGTGG TGGAT CCTGG AGATC G |
| reverse_primer_L+112 | Cy5-GCGAG ATTAC CATTA AGTGA A |
| reverse_primer_L+212 | Cy5-GACAA CTTTA CTGGG GCGAA CTAAA AC |
| reverse_primer_L+312 | Cy5-GCTTC CATAC CACCT GGCGC CATCC AT |
| reverse_primer_L+512 | Cy5-GGTCT AAGCT TGTGG TGTCT TGTGT GT |
| lambda_forward_primer | GTTTT CTGGG TTGGT |
| lambda_reverse_primer | GGCGG GTTTT GTTTT |
| forward_primer_extension | ACTAT CTATT CTCCC ATCTA TCAAA AAGAG TATTG ACTTA AAGTC |
| forward_primer_biotin_α | Biotin-ACTAT CTATT CTCCC ATC |

### Supplementary Figures

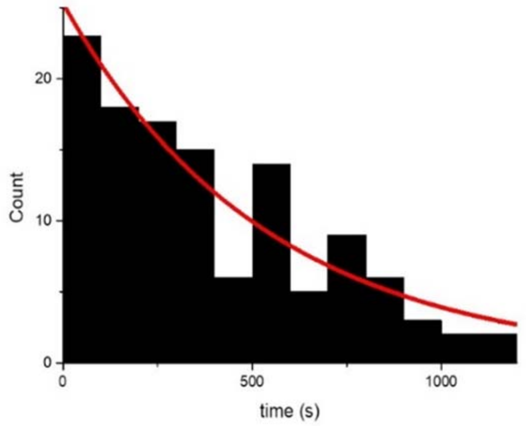


**Fig. S1. RNAP·DNA complex duration at the DNA end after RNA release.** We measured Cy5 PIFE survival time after the RNA release, and the distribution was fitted to a single exponential function to obtain the RNAP's retention time of 536±83 s.

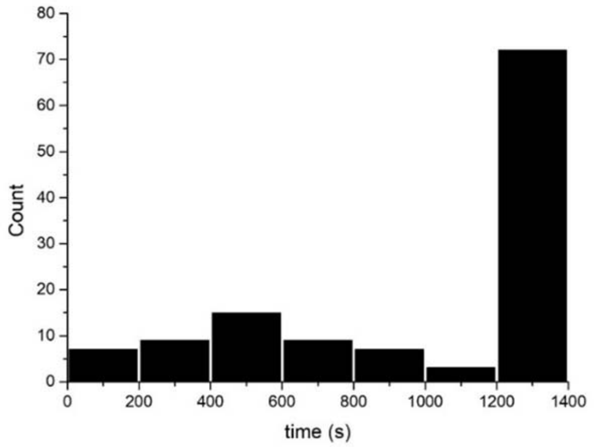


**Fig. S2. Cy5 photobleaching time.** To measure Cy5 survival time, we performed single-molecule imaging without NTP injection. About 60% of the molecules survived for longer than 1200 s after single-molecule imaging started, from which the photobleaching time of Cy5 was estimated as 2350 s.

**Fig. S3. Sigma retention.** We estimated the percentage of transcription complex containing $\sigma^{70}$ by comparing the number of Cy3-Cy5 colocalized spots between the cases of Cy5-end of DNA and those of Cy5-$\sigma$. The Cy5 labeling efficiencies of DNA (99%) and $\sigma^{70}$ (105%) were similar. When Cy5 was labeled at the DNA end, 46±10% of Cy3 signal was colocalized with Cy5. When Cy5 was labeled on $\sigma^{70}$, 35±6% of Cy3 signal was colocalized with Cy5. In the figures, the green circles, red dots, and black dots indicate all Cy3 spots identified, Cy3 spots colocalized with Cy5 signal, and the Cy3 spots non-colocalized with Cy5 signal, respectively. From these results, we estimated that 75% (34.5/45.8) of elongation complex has $\sigma^{70}$.

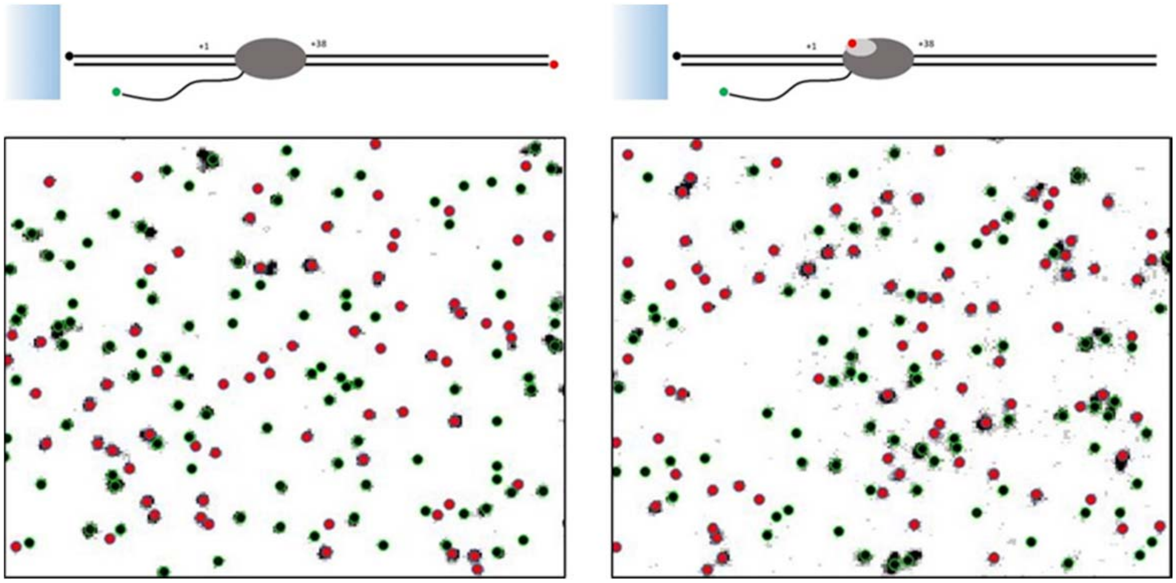

**Fig. S4. σ factor lasting time at the DNA end after RNA release.** We measured Cy5 signal vanishing time after NTP addition in termination complexes of Fig. 1F template.

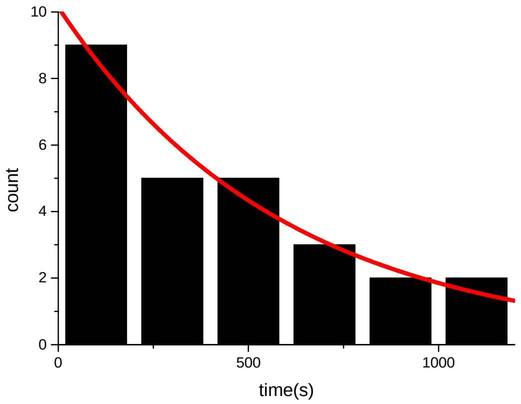

**Fig. S5. E111Q inhibits the cleavage of DNA by EcoRI.** Using the L+112 DNA template labeled with Cy5, we measured the cleavage inhibition efficiency of 1 nM E111Q. We counted the number of Cy5 spots before (left) and at 10 min after wild-type EcoRI injection (right) without (top) and with E111Q incubation for 5 min (bottom) before starting the imaging. The number of spots was decreased by 86±2% without E111Q incubation. On the other hand, the number was decreased by 26±4% with E111Q incubation. From these data, we estimated the cleavage inhibition efficiency of E111Q as 71±5%.

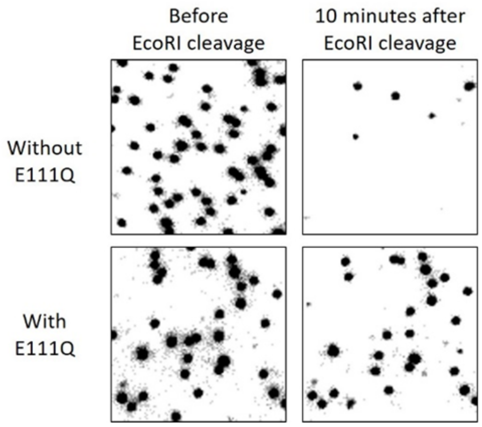


**Fig. S6. Diffusion time of RNAP on a long DNA template.** We prepared a transcription complex with Cy5-labled $\sigma^{70}$ on a 1560-bp DNA template (L+lambda), and measured the Cy5 signal vanishing time after termination.

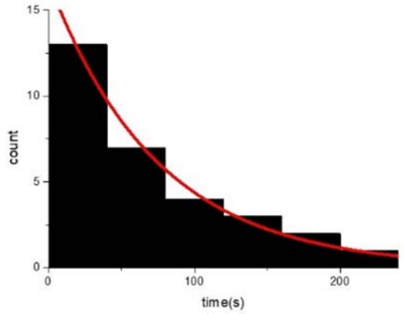


**Fig. S7. Termination time of various templates.** We measured the Cy3 signal vanishing time after NTP addition for L+15 (A), L+62 (B), L+112 (C), L+112R (D), L+212 (E), L+312 (F), L+512 (G) and L+112 (H) with E111Q. The distributions were fitted to a single-exponential decay function, and the fitted decay times are summarized on table S1.

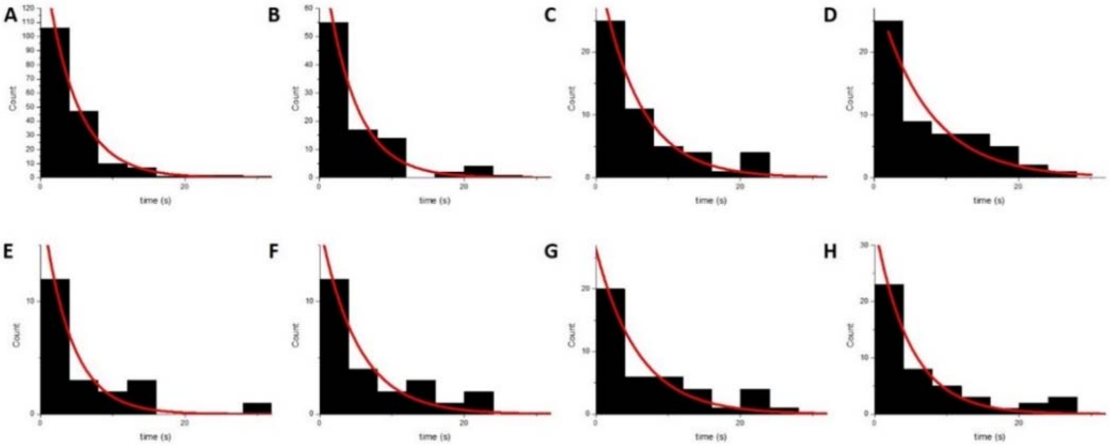
A
B
C
D
E
F
G
H
Count
time (s)
